# Human *Trypanosoma cruzi* infection risk is driven by eco-social interactions in rural communities of the Argentine Chaco

**DOI:** 10.1101/627141

**Authors:** Maria del Pilar Fernández, Maria Sol Gaspe, Paula Sartor, Ricardo E. Gürtler

**Affiliations:** Universidad de Buenos Aires. Departamento de Ecología, Genética y Evolución, Facultad de Ciencias Exactas y Naturales, Ciudad Universitaria, C1428EHA, Buenos Aires, Argentina; Consejo Nacional de Investigaciones Científicas y Técnicas. Instituto de Ecología, Genética y Evolución de Buenos Aires (IEGEBA), Ciudad Universitaria, C1428EHA, Buenos Aires, Argentina; Earth Institute, Columbia University. New York, NY, 10025, United States of America; Ministerio de Salud Pública del Chaco H3500ACC, Resistencia, Chaco, Argentina; Facultad de Ciencias Exactas, Naturales y Agrimensura, Universidad Nacional del Nordeste W3404AAR, Corrientes, Corrientes

**Author notes:** Corresponding author María del Pilar Fernandez.

## Abstract

The transmission of *Trypanosoma cruzi* to humans is determined by multiple ecological, socio-economic and cultural factors acting at different scales. Their effects on the human risk of infection with *T. cruzi* have often been examined separately or using a limited set of ecological and socio-demographic variables. Herein, we integrated the ecological and social dimensions of human disease risk with the spatial distribution patterns of human and vector (*Triatoma infestans*) infection with *T. cruzi* in rural communities of the Argentine Chaco composed mainly of indigenous people (90% Qom) and a creole minority. Prior to the implementation of a vector control intervention, the estimated seroprevalence of *T. cruzi* among 1,929 local residents examined in a cross-sectional study was 29.0%, and was twice as large in Qom than creoles. Using generalized linear mixed models, the risk of human infection increased by 60% with each additional infected triatomine and by 40% with each seropositive household co-inhabitant; increased significantly with increasing household social vulnerability (a multidimensional index of poverty), and decreased with increasing host availability in sleeping quarters. A significant negative interaction between household social vulnerability and the relative abundance of infected *T. infestans* indicated that vulnerable household residents were exposed to a higher risk of infection even at low infected-vector abundances. Household mobility within the study area reduced the effects of domiciliary vector abundance, possibly due to less consistent exposures. Nonetheless, the seroprevalence rates of movers and non-movers were not significantly different. Human infection was clustered by household and at a larger spatial scale, with hotspots of human and vector infection matching areas of higher social vulnerability. These results were integrated in a risk map that shows high-priority areas for targeted interventions oriented to suppress house (re)infestations, treat infected children, and thus reduce the burden of future disease.

**Author summary:** Chagas disease is one of the main neglected tropical diseases (NTDs) affecting vulnerable communities in Latin America where transmission by triatomine vectors still occurs. Access to diagnosis and treatment is one of the remaining challenges for sustainable control of Chagas disease in endemic areas. In this study, we integrated the ecological and social determinants of human infection with the spatial component to identify individuals, households and geographic sectors at higher risk of infection. We found that the risk of human infection was higher in indigenous people compared to creoles, and increased with the abundance of infected vectors and with household social vulnerability (a multidimensional index of poverty). We also found that the social factors modulated the effect of the abundance of infected vectors: vulnerable-household residents were exposed to a higher risk of infection even at low infected-vector abundance, and human mobility within the area determined a lower and more variable exposure to the vector over time. These results were integrated in a risk map that showed high-priority areas, which can be used in designing cost-effective serological screening strategies adapted to resource-constrained areas.

## Introduction

The transmission of zoonotic and vector-borne diseases is an “inherently ecological process” involving intraspecific and interspecific interactions between vector, pathogen and host populations [1]. As shown by Ross-Macdonald style mathematical models, the transmission of vector-borne pathogens is determined by the density and survival of insect vectors, vector-host contact rate, host susceptibility, vector and host infectivity, and duration of infection [2,3]. When human populations are implicated, pathogen transmission dynamics also involve socio-economic, cultural, political, psychological and ethical factors, which pertain to the human dimension of disease [4,5]. With a focus on human health, these ecological, biological and social factors affecting the transmission of vector-borne diseases may be classified as intrinsic or extrinsic to the human population [6]. Intrinsic factors are biological in nature (i.e., immune response) and can only be manipulated by advances in medical research and technology [6]. Extrinsic factors include the environmental context, vector ecology and behavior, human activities, socio-economic inequalities and political factors, among other [6]. A thorough understanding of the combined effects of the extrinsic factors is required to design more effective and sustainable disease control interventions, which have traditionally been crafted in a reductionist biomedical approach [7]. Reductionism argues that the sum of information provided by studying each component of the system separately is sufficient to understand disease transmission dynamics [8]. By contrast, the Ecohealth or eco-bio-social approaches focus on the interactions among multiple ecological and social factors and their combined effects on human health as part of a complex socio-ecological system [7].

Chagas disease is a chronic infection caused by the protozoan *Trypanosoma cruzi*, and is mainly transmitted by triatomine vectors in endemic areas [9] in spite of regional and intergovernmental vector control (and elimination) initiatives, such as the INCOSUR (“Iniciativa de Salud del Cono Sur”) launched in the 1990s in the Southern Cone countries in South America [10]. Recent estimates indicate that 62.4% of people infected with *T. cruzi* live in INCOSUR countries, and 30.6% of new cases due to vector-borne transmission occur in Bolivia and Argentina [9], where the Gran Chaco region remains a hotspot for Chagas disease [11,12]. Although in the Argentine Chaco region the overall human seroprevalence of *T. cruzi* has declined over the last 60 years [13–15], it remains high (27.8-71.1%) in rural communities encompassing creole and indigenous populations [16–24].

In the Gran Chaco, human infection with *T. cruzi* usually occurs within sleeping quarters before reaching 15 years of age and is transmitted by *Triatoma infestans* [25–27]. Most studies of human infection with *T. cruzi* in endemic areas have focused on the seroprevalence distribution among demographic subgroups and/or on the effects of vector presence, abundance and *T. cruzi* infection status. Although the association between human *T. cruzi* infection and selected socio-demographic factors has been investigated [24,25,28–31], these studies either did not address the combined effects of ecological and social variables due to limited data availability, or only considered a few socio-demographic variables. Human infection was positively associated with the presence or abundance of domestic animals [24,25,28,29,32]. A less clear association was found between *T. cruzi* human infection and house construction quality (i.e., thatched roofs and cracks in the walls): while in some studies human infection increased in poor-quality housing [25,28,31,32], others did not find such association [29,30].

The joint analysis of the spatial distribution of human and vector infection can shed light into the processes and factors associated with the vector-borne transmission of *T. cruzi*. Integrating the spatial component of disease with household-level and individual-based risk factor analysis is needed to identify transmission hotspots, create risk maps of *T. cruzi* infection, and stratify the affected areas for targeted control [33–35]. For Chagas disease, spatial analysis has been used to investigate the reinfestation process with *T. infestans* after community-wide insecticide spraying [36–40]; identify factors associated with house infestation [41–43]; evaluate the effects of vector control interventions [44], and generate risk maps at regional and continental levels based on vector distribution [45,46]. By contrast, the spatial component of human *T. cruzi* infection has rarely been investigated, and the ones that did it focused on periurban settings [29,47,48].

As part of a broader research and control program on the transmission of *T. cruzi* in rural communities of the Argentine Chaco [24,40–42,49–53], this study investigated the effects of household-level vector indices and demographic factors on human *T. cruzi* infection at individual, household and community levels. We also considered the household socio-economic status and the interaction between social and ecological variables using indices of social vulnerability (as a measure of socio-economic inequalities) and of domestic host availability, i.e., in human sleeping quarters or domicile [54]. Both indices were positively associated to the abundance of infected domiciliary vectors [54]. Herein, we hypothesize that the interaction between ecological and social factors would enhance the risk of *T. cruzi* transmission to humans, which cannot solely be explained by variation in vector indices. At a community level, we evaluated the spatial correlation between human and vector infection, host availability and social vulnerability. We integrated the evidence provided by the individual-based risk model with the spatial component to derive community-level risk maps of human infection, with an emphasis in children, which represent the target demographic group for *T. cruzi* infection diagnosis and treatment.

## Methods

### Study area

This study was conducted in a rural section of Pampa del Indio municipality (25° 55’S 56° 58’W), Chaco province, Argentina, denominated Area III, which encompassed three contiguous large communities (Cuarta Legua, Pampa Grande and Pampa Chica), and other smaller ones [41]. This area was subjected to a vector control and Chagas disease research program initiated in 2008 with a follow-up period of 7 years as of 2015. In October 2008, a baseline vector survey found that a third of the inhabited houses were infested with *T. infestans*, mainly within human sleeping quarters, and virtually all (93.4%) houses were sprayed with insecticides immediately after [41]. During the 2008-2015 vector surveillance phase we conducted annual triatomine surveys and selectively sprayed with insecticide the few foci detected. This strategy reduced house infestation to <1% during 2008-2012, and no infested house was found in 2015 [40]. Prior to 2008, vector control actions had been sporadic and partial; the last community-wide insecticide spraying campaign took place in 1997-1998, and the last sprays registered prior to the baseline survey were carried out in a few houses in 2006 and July 2008 [41].

The population in the area monotonically increased during the 2008-2015 period, from 2,392 people and 407 inhabited houses in 2008 to 2,548 people and 587 houses in 2015 [54]. The population was composed of two ethnic groups; 93.1% of the residents self-identified as Qom people and the remaining minority as creoles [54]. A household was defined as all the people who occupy a housing unit, including related and nonrelated family members [55]. The housing unit (or house compound) included one or more domiciles (i.e., sleeping quarters) and associated peridomestic structures.

### Study design

This study complied with STROBE recommendations for observational studies [56] (S1 Text). We integrated vector and epidemiological information collected during the 2008-2015 period during the annual vector and household surveys as described in detail elsewhere [40,41,54]. At each visit, the location of the domicile was georeferenced with a GPS receiver (Garmin Legend) and the demographic information of the household updated. The household surveys included relevant epidemiological data such as number, type and construction materials of the different sites in each housing unit, refuge availability for the vector, the number and resting places of domestic animals, number of house occupants, and their ages and ethnicity. In 2012-2015, we carried out a thorough census of the population, which included several social variables identified as social determinants of health and relevant to NTD transmission, such as educational level, income source, household mobility, access to health services, etc [54]. Response rates were high (>90%) in all surveys [40].

Four years after community-wide insecticide spraying and with a low risk of vector-borne transmission (<1% of infested houses), we conducted a series of serosurveys aimed at total population coverage (older than 9 months old) during the 2012-2015 period. Using a participatory approach, we had meetings with the local population to discuss project tasks and coordinate activities with local healthcare agents and hospital personnel [50]. During 2012-2013, people were recruited passively through the radio, schools and word of mouth; venipuncture was done at schools and at the primary health posts of each large community to draw a blood sample of 3-7 ml. In April 2015, recruitment was done actively and blood draws were conducted coupled to the vector survey. The protocol is described in detail elsewhere [24,50].

### Ethics statement

This study complied with the ethical principles included in the Declaration of Helsinki (Ethical Committee “Dr. Carlos A. Barclay”, Protocol ref. TW-01-004). All adult subjects provided written informed consent prior to participation in the serosurvey, and a parent or guardian of any child participant younger than 14 years old provided informed consent on the child’s behalf. Participants aged between 14 and 18 years old, provided their written informed consent along with consent from a parent or guardian. Informed consent forms were provided in Spanish and translated to Qom language (*Qomlactaq*) orally by primary healthcare agents when needed. Oral consent was obtained for household surveys, including entomological searches, as approved by the IRB and recorded in the data collection sheets as described [41].

### Serodiagnosis of human infection

Blood samples were transported to the local hospital where serum samples were obtained by centrifugation at 3,000 rpm for 15 min within the same day. Each serum was allocated in triplicate vials and stored at −20°C. Serum samples were tested using two different ELISA tests based on conventional (Chagatest, Wiener) and recombinant antigens (ELISA Rec V3.0, Wiener) as described elsewhere [50]. Discordant results between both ELISAs were resolved via an indirect immunofluorescence antibody test conducted at the reference center for Chagas disease serodiagnosis (Instituto Nacional de Parasitología Dr M. Fatala Chaben). A person was considered seropositive (i.e., infected) if at least two tests were reactive. The serosurvey results were informed to the participants within 8 weeks of the blood draw, and follow-up of *T. cruzi*-seropositive people was done by physicians from the local hospital as described elsewhere [50], including treatment with benznidazole or nifurtimox of people younger than <18 y.o. and those adults who specifically requested it.

### Vector-related indices

All triatomines collected at the baseline vector survey in 2008 were identified taxonomically and the individual infection status with *T. cruzi* was determined by microscope examination of feces [41] or by molecular diagnosis using kDNA-PCR [57], achieving a coverage of 60% of all infested houses. The occurrence of domiciliary infestation with *T. infestans* was determined by the finding of at least one live triatomine (excluding eggs) through any of the vector collection methods used (i.e., timed-manual searches, bug collections during insecticide spraying operations, and householders’ bug collections). The relative abundance of domiciliary *T. infestans* was calculated only for infested houses as the number of live bugs collected by timed-manual searches per 15 min-person per site, as described [41]. The same procedures were used to determine the occurrence and abundance of *T. cruzi*-infected *T. infestans* in the domicile.

### Epidemiological data pre-processing

Human seropositivity data (collected over 2012-2015) were back-corrected to 2008 in order to analyze the risk of human infection prior to the community-wide insecticide spraying campaign when vector-borne transmission was still active, and given the virtual absence of house infestation and of infected triatomines over 2008-2012. This allowed us to compute a more accurate length of exposure to vector-borne infections by subtracting the time elapsed since the baseline vector survey to the age recorded at the serosurvey. Using the demographic data collected in household surveys in 2008-2015 and the location of each household over the different time periods, we identified people’s residential location as of 2008. The 2012-2015 detailed census data was also used to determine their mobility pattern, which was classified as: non-movers (individuals whose house remained at the same geographical location), movers (if they moved within the study area), or as in-migrants or out-migrants (to or from outside the study area, respectively, including individuals coming from or leaving to a different section within Pampa del Indio municipality), as described elsewhere [54]. If the in-migrants had resided in the study area prior to 2008, we classified them as return migrants.

Regarding the socio-demographic variables collected in 2012, they were back-corrected to 2008 whenever possible, assuming similar socio-economic conditions prevailed over 2012 and 2008 [41]. Data obtained from the household surveys were used to construct two surrogate indices (the social vulnerability index and the host availability index) by means of multiple correspondence analysis. A detailed description of the indices, their rational and construction is found elsewhere [54]. Briefly, the social vulnerability index estimated for 2008 households included characteristics of the domiciles (refuge availability, presence of cardboard roofs and/or mud walls, time since house construction, and domestic area), and household socio-economic and demographic characteristics (overcrowding, goat-equivalent index and educational level). This index aimed at summarizing the multiple interrelated socio-demographic variables associated with socio-economic position. The host availability index in domiciles summarized the number of potential domestic hosts of *T. infestans* (i.e., adult and child residents, total number of dogs, cats, and chickens nesting indoors), and in the case of dogs and cats, whether they rested within or in the proximity of the domicile. A preliminary analysis showed that the household abundance of domestic animal hosts was positively correlated with larger household size, i.e., number of human residents [54].

### Data analysis

For each variable we checked whether the missing values were missing completely at random by building a dummy variable (missing and non-missing values) and analyzing the significance of Spearman’s correlation coefficient with any another independent variable in the data set, as described elsewhere [41]. Most of the variables with missing values (collected through household surveys) were missing completely at random, except for educational level and overcrowding in 2008, in which the missing data corresponded to households that had moved or out-migrated by 2012 (the year when these data were collected). Missing data for human infection per household were also biased towards adult males (who refused to participate more often than other demographic groups), and movers, as we were not able to assign their residential location in 2008 to all of them. We excluded from the analysis 197 people tested who resided elsewhere but were visiting the study area at the time of the serosurvey. Throughout the text, minors younger than 15 y.o. are referred to as children for simplicity.

For all proportions, 95% confidence intervals (CI_95_) were estimated using the Agresti and Coull method if sample sizes were greater than 50, and the Wilson method for smaller sample sizes [58]. We used χ^2^ tests for bivariate analysis of categorical variables; generalized linear models (GLM) [59] with logit link function when the outcome of interest had a binary distribution, and a negative binomial GLM (link function: log) when analyzing count data, such as the number of seropositive people. In the case of binary response variables, the relative risk was expressed as Odds Ratios (OR), and in the case of count data as incidence rate ratios (IRR). Negative binomial regression was preferred to Poisson regression given the overdispersed distributions [60]. Mixed-effects models (GLMM) were also considered when individual data was analyzed to account for possible household-related random effects [24,28]. All analysis were implemented in Stata 14.2 [61] and R 3.2.3 (lme4 and car packages) [62].

### Infection risk models

The infection risk model was estimated for the entire population and for children separately. The outcome of interest was their seropositivity status for *T. cruzi* (yes/no). In the model for the total population we included age, gender, ethnicity, and household-level variables: the relative abundance of *T. cruzi*-infected *T. infestans* per unit of catch-effort in the domicile, the social vulnerability index, host availability index, and the number of seropositive co-inhabitants. The model for child infection also included the mother’s seropositivity status, and whether any recent insecticide spraying occurred in 2006-2008 prior to the baseline vector survey. We also evaluated the interaction between vector abundance, the social vulnerability and host availability indices, and retained in the model the ones that had a significant effect. We compared a GLMM model (logit link function) considering the household as a random variable and a GLM model in both cases.

We used an information theoretic approach and Akaike’s information criterion (AIC) to identify the best-fitting models given the data collected, and a multimodel inference approach to account for model selection uncertainty [47,48] using the MuMin R-package. This approach was used to identify the ecological and social factors associated with house infestation, as described elsewhere [41]. Odds Ratios (ORs) and their 95% confidence intervals were calculated from model-averaged coefficients. The relative importance (RI) of each variable is defined as the sum of Akaike weights in each model in which the variable is present; RI takes values from 0 to 1. Multicollinearity was assessed by the Variance Inflation Factor (VIF), and model fitting for the logistic regression was assessed by the Hosmer-Lemeshow goodness of fit test and the Receiver Operator Curve (ROC), including the area under the ROC (AUC) [62]. To assess sensitivity and specificity of the models, we employed an optimal threshold value that minimized the sum of error frequencies [63]. This value was obtained by finding the maximum sum of sensitivity and specificity for all threshold values t (sens(t)+spec(t)) using the pROC R-package [64]. The multimodel inference approach does not allow for missing data, therefore individuals with missing data were not considered in the models.

### Spatial analysis and risk maps

Global point pattern analysis (univariate and bivariate) were performed for human and vector infection using the weighted K-function implemented in Programita [35]. Random labeling was used to test the null hypothesis of random occurrence of events among the fixed spatial distribution of all households. We used quantitative and qualitative labels for each household as previously described [54]. Monte Carlo simulations (n = 999) were performed and the 95% ‘confidence envelope’ was calculated with the 2.5% upper and lower simulations. Additionally, local spatial analysis on the abundance of (infected) vectors were performed using the G* statistic implemented in PPA [33]. The selected cell size was 200 m (assuming that each household had at least three neighbors at the minimum distance of analysis), and the maximum distance was set at 6 km (i.e., half of the dimension of the area). We created heatmaps (i.e., density maps) to visualize the spatial aggregation of the variables using a kernel density estimation algorithm within a radius of 200 m as implemented in QGIS 2.18.11. Risk maps were obtained by interpolating the predicted probabilities from the risk model and using the 2008 household location. The geographical coordinates of each housing unit were transformed to preserve the privacy of the households involved in this study, as described elsewhere [41].

## Results

During 2012 and 2015, we tested 1,929 people that resided in the study area at the time of the serosurveys. Passive recruitment during 2012-2013 at local schools and primary health posts led to a coverage of 47.8% of the local population, which increased to 77.0% during 2015, when active recruitment of participants was coupled with vector surveys (S1 Fig). Diagnosis coverage was highest in school children (80.9%) and adult women (76.8%) in general (Fig 1).

**Fig 1.**
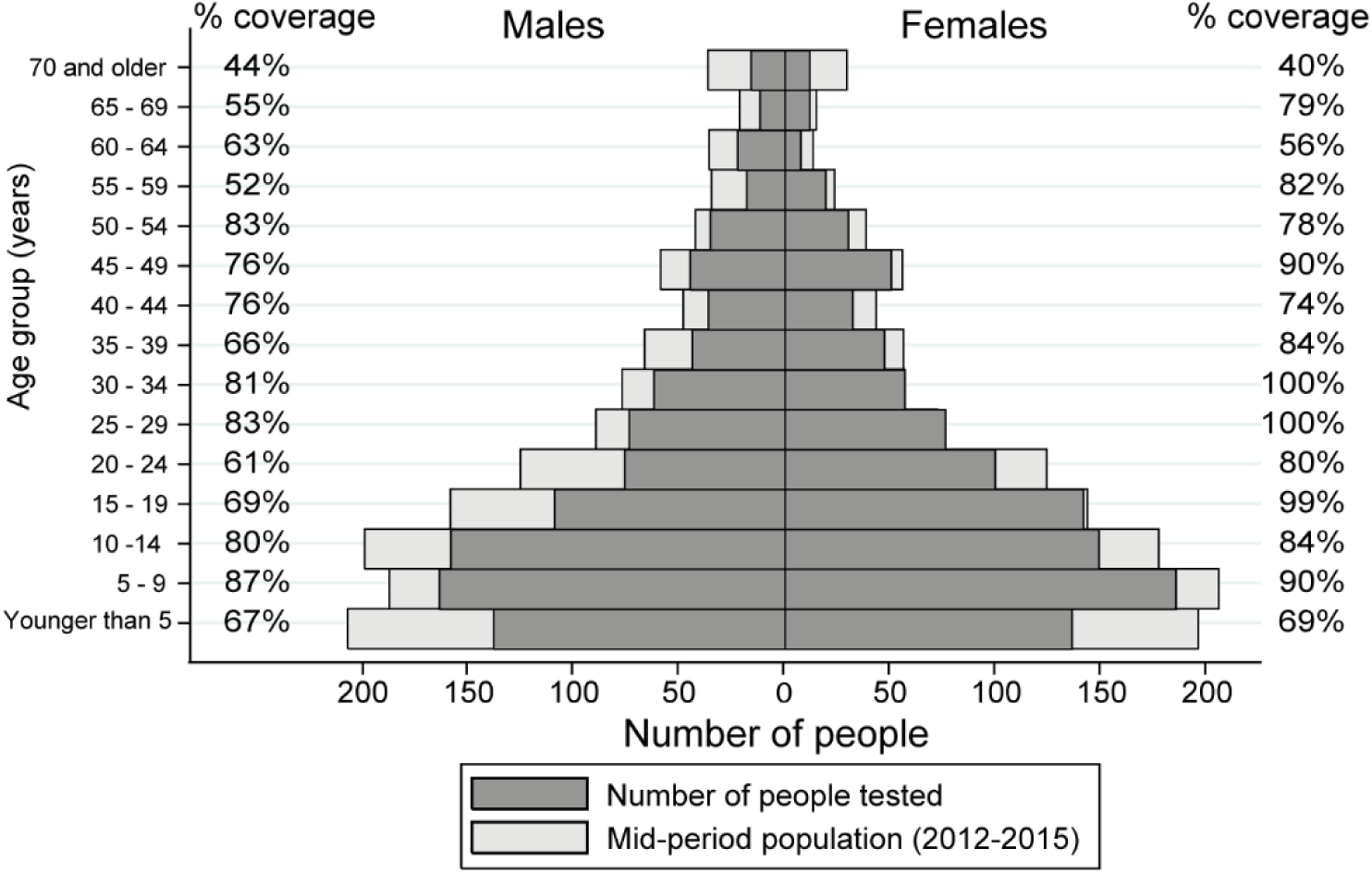
The 2012-2015 mid-period total population per 5-year age group and the population serologically tested for *T. cruzi* infection (number and percentage) in Pampa del Indio, Argentina.

The two-tiered ELISA serological testing showed almost perfect agreement between tests (kappa index = 0.9, p < 0.001). Seropositive results for *T. cruzi* infection were observed in 25.3% of the samples, whereas 5.5% that were discordant later tested negative at the reference laboratory. Only 1.5% (CI_95_ = 4*10^-4^-3.0%) of children born after the community-wide insecticide spray in 2008 were seropositive. All four *T. cruzi*-seropositive children born post-intervention had *T. cruzi*-seropositive mothers and no vectors were collected in their domiciles during the surveillance phase. For the same age group (children <6 y.o.), the risk of infection was higher if they had been born before the community-wide spraying (logistic regression, OR_2008_ = 6.2, CI_95_ = 1.2-31.8, p = 0.01), after adjusting for age. Only 1.4% of the people tested had been exposed to *T. infestans* due to transient domiciliary reinfestations during the surveillance period, but all 38 also lived in infested houses prior to the 2008 insecticide campaign.

We were able to assign the local residency status as of 2008 (i.e., their house identification code in 2008) to 82.6% of the people tested. The overall seroprevalence estimated for 2008 was 29.0% (CI_95_ = 26.7-31.4, n = 1373). It increased significantly with age from 5.7% in children younger than 5 y.o. to 25% in teenagers (14 to 19 y.o.), then jumped to 50% in young adults (20 to 29 y.o.), and remained around 60% in older adults (Fig 2). Although females had a lower overall seroprevalence rate than males (26.5 vs 31.6%; χ^2^ test, df =1, p = 0.04), this difference was more evident in adults (Fig 2) and was not significant after adjusting for age (S1 Table). The seroprevalence for Qom people almost doubled that observed for creoles (29.7 vs 18.7%; χ^2^ test, df = 1, p = 0.02) (S1 Table).

**Fig 2.**
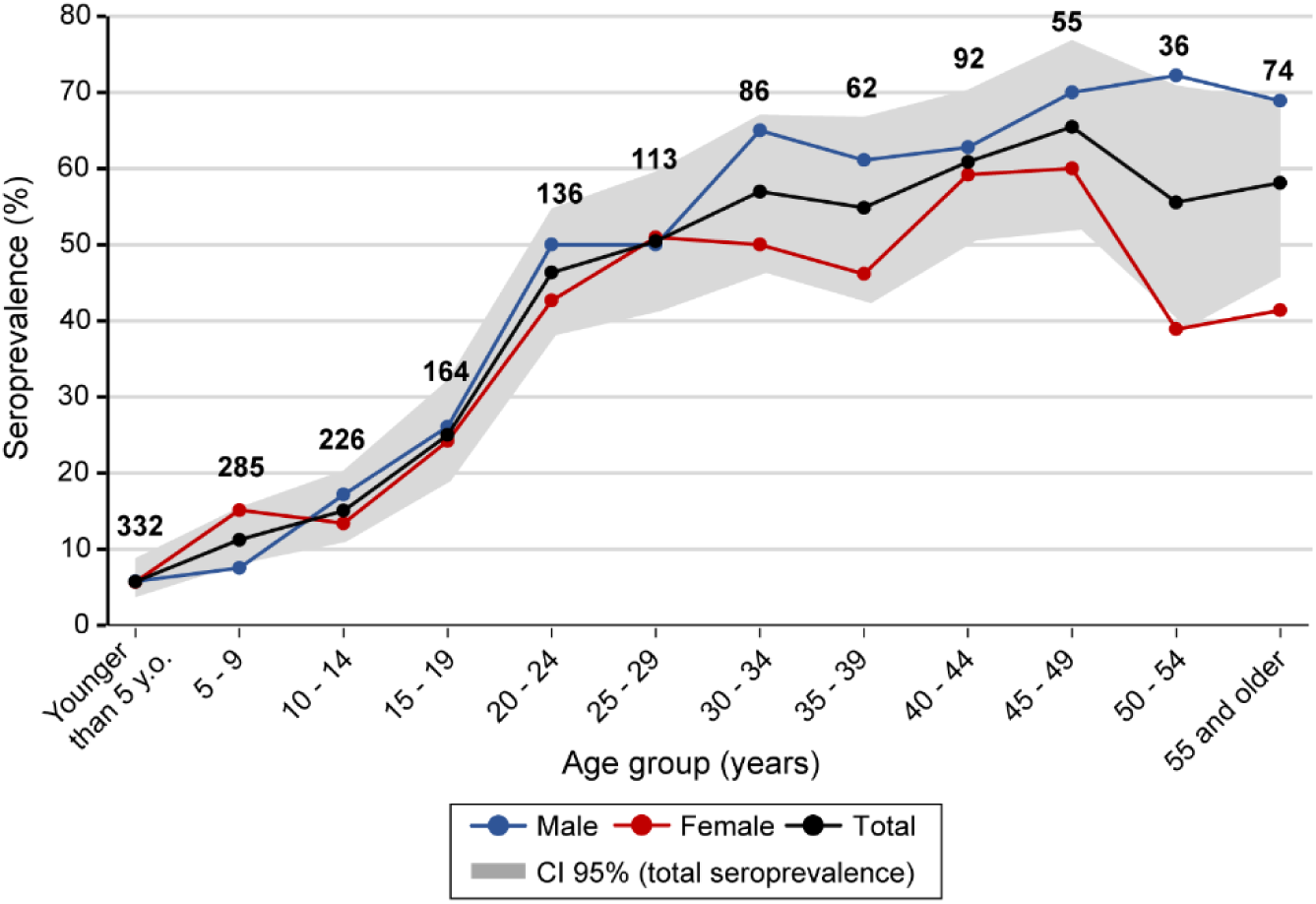
Seroprevalence of *T. cruzi* infection per 5-year age group and gender for the population in 2008, prior to a community-wide insecticide spraying campaign, Pampa del Indio, Argentina. The numbers above the lines indicate the total number of people serologically tested.

### Household infection and domiciliary infestation

We found at least one *T. cruzi*-seropositive person in 71.3% of the 301 households in which at least one member had been tested (77% of all households existing in 2008). For children, that proportion dropped to 16.7%. The occurrence of at least one seropositive person per household was not associated with the abundance of *T. cruzi*-infected *T. infestans* per unit effort in the domicile (logistic regression, OR =1.5, CI_95_ = 0.8-3.0; p = 0.2). In contrast, the occurrence of at least one infected child was positively and highly significantly associated with the abundance of infected vectors (logistic regression, OR = 2.8; CI_95_ = 1.6-4.9; p < 0.001). The household number of seropositive people increased with the abundance of infected vectors (negative binomial regression, IRR = 1.1; CI_95_ = 1.1-1.2; p < 0.001).

The seroprevalence of *T. cruzi* was higher in local residents who reported past exposure to *T. infestans* at the time of the serosurvey (35.3 vs 15.0%, χ^2^ test, df = 1, p < 0.001). There was a fair agreement between the house infestation status assessed in 2008 and reported past exposure to triatomines (kappa index = 0.2; p < 0.001). Although 79% of residents in an infested house reported past exposure to the vector, a considerable proportion (54%) of those who inhabited non-infested houses also reported past exposures. Therefore, the individual infection risk adjusted for age, gender and ethnic group was two-fold higher in infested houses as determined in the 2008 (baseline) vector survey (GLMM, OR = 2.3, CI_95_ = 1.4-3.9, p < 0.001), but the effect was marginally significant when the reported past exposure was considered instead (GLMM, OR = 1.5, CI_95_ = 1-1.2, p = 0.6) (S2 Table). In children, the infection risk in infested houses was almost four-fold higher (GLMM, OR = 3.7, CI_95_ = 1.3-10, p = 0.01) even after adjusting for the mother’s seropositivity status, while the reported past exposure had no significant effect (GLMM, OR = 0.9, CI_95_ = 0.4-2.4, p = 0.9) (S2 Table). When vector abundance per unit effort was considered, the infection risk in children increased 42% per 5 additional *T. infestans* collected in the domicile (GLMM, OR = 1.42, IC_95_ = 1.2-1.7, p = 0.001).

### Household mobility and human infection

Movers (as registered in 2012-2015) had a 30-110% higher risk of residing in a *T. infestans*-positive house in 2008 than non-movers after adjusting for age, gender and ethnic group (logistic regression, OR = 1.7; CI_95_ = 1.3-2.1; p < 0.001). Out-migrants had a much lower risk (OR = 0.5; CI_95_ = 0.3-1.3; p < 0.01). These findings proved valid in children, whose infection status is more closely associated with recent domiciliary infestation. Although child infection was not directly associated with their mobility pattern, the effect of domiciliary infestation on infection risk was modified by mobility: in children living in infested houses in 2008, the effect of infestation on risk decreased by 90% in movers compared to non-movers (Table 1). While non-mover children residing in infested houses in 2008 had a six-fold higher risk of infection compared to non-mover children living in non-infested houses (20.0% vs. 4.5%, respectively) (Table 1), mover children living in infested houses had a nearly equal seroprevalence as those living in non-infested houses (11.9% vs. 11.1%) (Fig 3).

**Table 1.** Generalized linear mixed model of child seropositivity for *T. cruzi* infection in 2008 clustered by household (logit link function), Pampa del Indio, Argentina. The odds ratio (OR) of the interaction terms indicate if the effect of infestation for non-movers increased (>1) or decreased (<1) in other mobility categories (out-migrants or return migrants).

| Variables | OR (CI<sub>95</sub>) | P |
| --- | --- | --- |
| **Infestation (non-movers)** |  |  |
| No | 1 |  |
| Yes | 6.1 (1.9-19.0) | 0.002* |
| **Age** | 1.14 (1.03-1.3) | 0.01* |
| **Gender** |  |  |
| Male | 1 |  |
| Female | 1.5 (0.7-3.2) | 0.3 |
| **Ethnic group** |  |  |
| Creole | 1 |  |
| Qom | 7.1 (0.4-139.0) | 0.2 |
| **Mother infection status** |  |  |
| No | 1 |  |
| Yes | 8.0 (2.5-25.0) | <0.001** |
| **Mobility (2012-2015)** |  |  |
| Non-movers (non-infested houses) | 1 |  |
| Movers (non-infested houses) | 1.8 (0.4-8.5) | 0.5 |
| Out-migrants (non-infested houses) | 2.2 (0.2-20.0) | 0.5 |
| Return migrants (non-infested houses) | 0 | 1 |
| **Interactions§** |  |  |
| Movers*infested houses | 0.10 (0.01-0.99) | 0.05* |
| Out-migrants*infested houses | 0 | 1 |
| Return migrants*infested houses | 0 | 1 |

§ The reference category is non-movers living in infested houses
OR: Odds ratio; CI<sub>95</sub>: 95% confidence interval
\*\* $p < 0.001$ ; \* $0.001 \leq p \leq 0.05$ ; $\sim 0.5 < p < 1$

**Fig 3.**
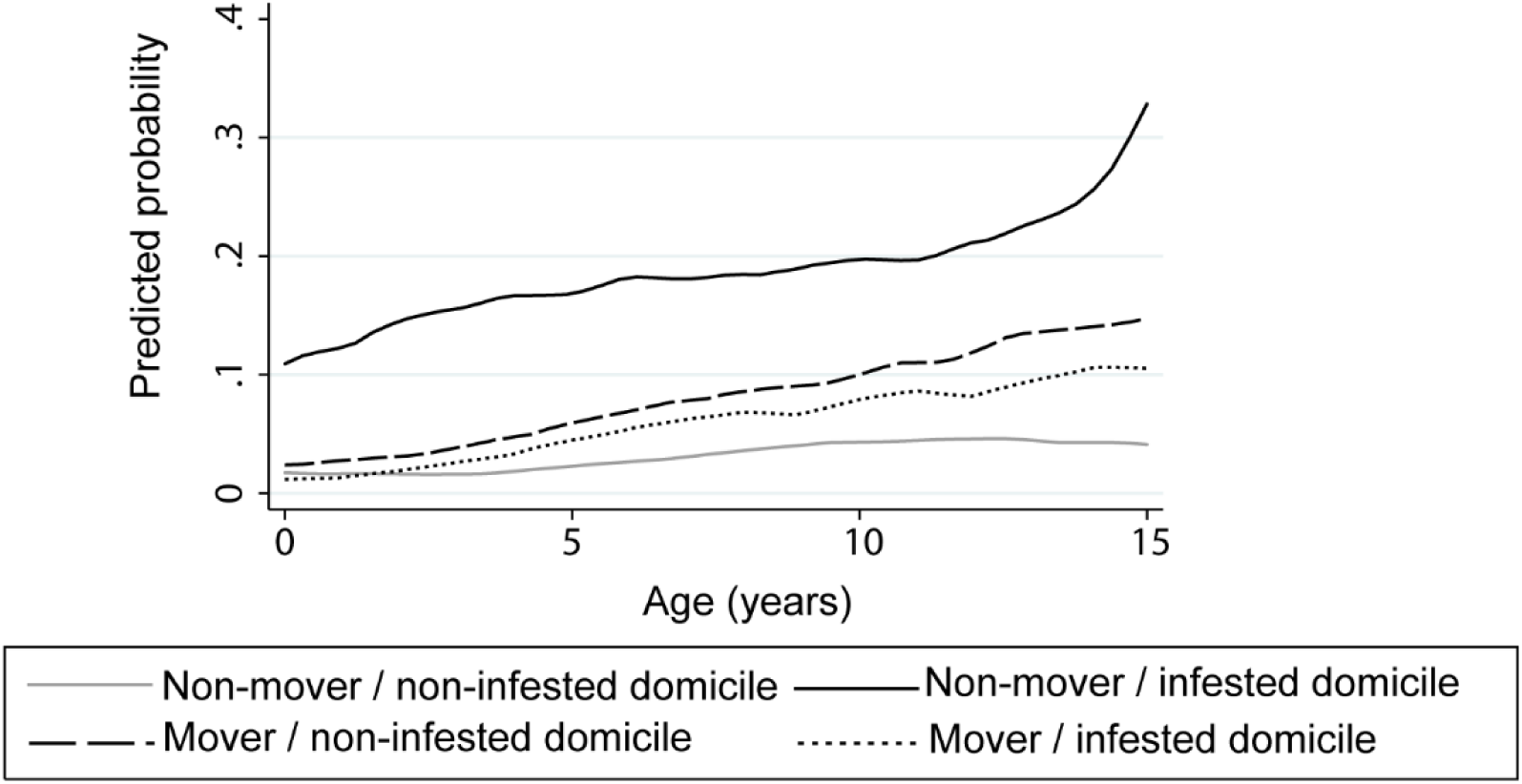
Predicted probabilities for the GLMM model of child seropositivity for *T. cruzi* infection regarding the interaction between domiciliary infestation and mobility patterns (non-movers vs. movers), Pampa del Indio, Argentina.

### Household socio-demographic determinants of human infection

The occurrence of at least one seropositive person per household was positively associated with social vulnerability (OR = 1.5; CI_95_ = 1.2-1.8; p = 0.001) and was independent of host availability (OR = 1.3; CI_95_ = 0.9-1.7; p = 0.1). The household presence of at least one seropositive child occurred more frequently in houses with high host availability (OR = 2.1; CI_95_ = 1.4-3.4; p < 0.001) and high social vulnerability (OR = 1.4; CI_95_ = 1.0-2.0; p = 0.05) (Fig 3). In both cases, the effects of the socio-demographic variables were additive, as the interaction terms were not significant. The occurrence and abundance of *T. cruzi*-seropositive children were similar to the distribution of infected domiciliary *T. infestans*, which also increased with host availability and social vulnerability (Fig 4).

**Fig 4.**
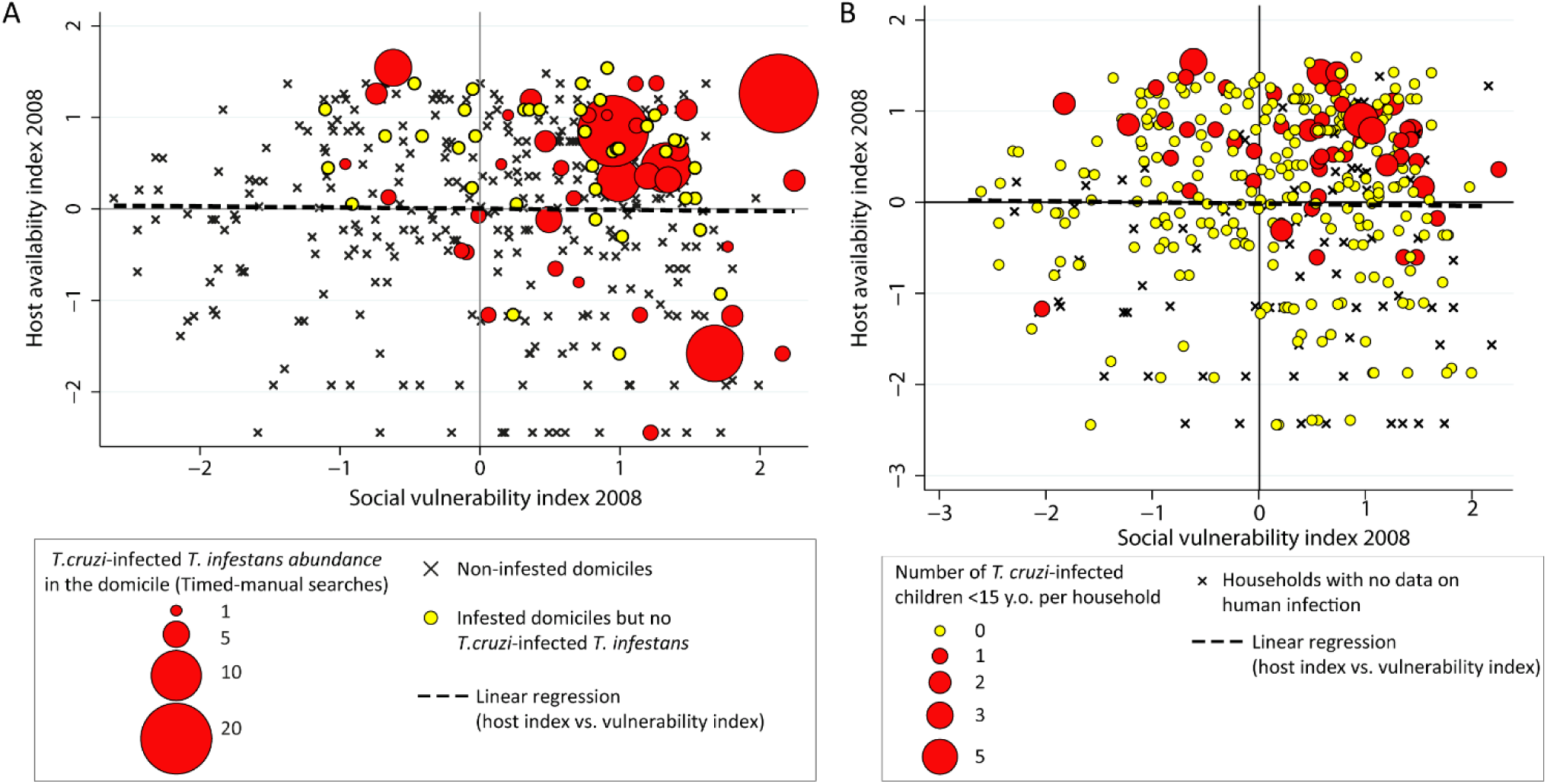
Multivariate relationship between household-level host availability, social vulnerability and vector indices [54] **(A)** and child seropositivity for *T. cruzi* infection **(B)**, Pampa del Indio, Argentina.

### Spatial distribution of human infection and house infestation

Despite the widespread occurrence of at least one *T. cruzi*-seropositive person per household, human infection was spatially aggregated in 2008 (Fig 5A). The global spatial analysis indicated that households with at least one seropositive person were aggregated at a scale between 2 and 6 km (S2A Fig), implying that up to 2 km (i.e., within each community) the spatial distribution was not significantly different from a random pattern. This result is consistent with the differences among communities in the proportion of households with at least one seropositive person: 84.6% in Cuarta Legua, 69.4% in Pampa Grande and 56.8% in Pampa Chica (χ^2^ test, df = 2, p < 0.001). When the number of seropositive people per household was considered, we did not find a significant spatial correlation between neighboring houses at any scale. Although the trend shown by the variograms indicated similar numbers of seropositive people between neighboring households, this pattern was not significantly different from a random distribution (S2B Fig).

**Fig 5.**
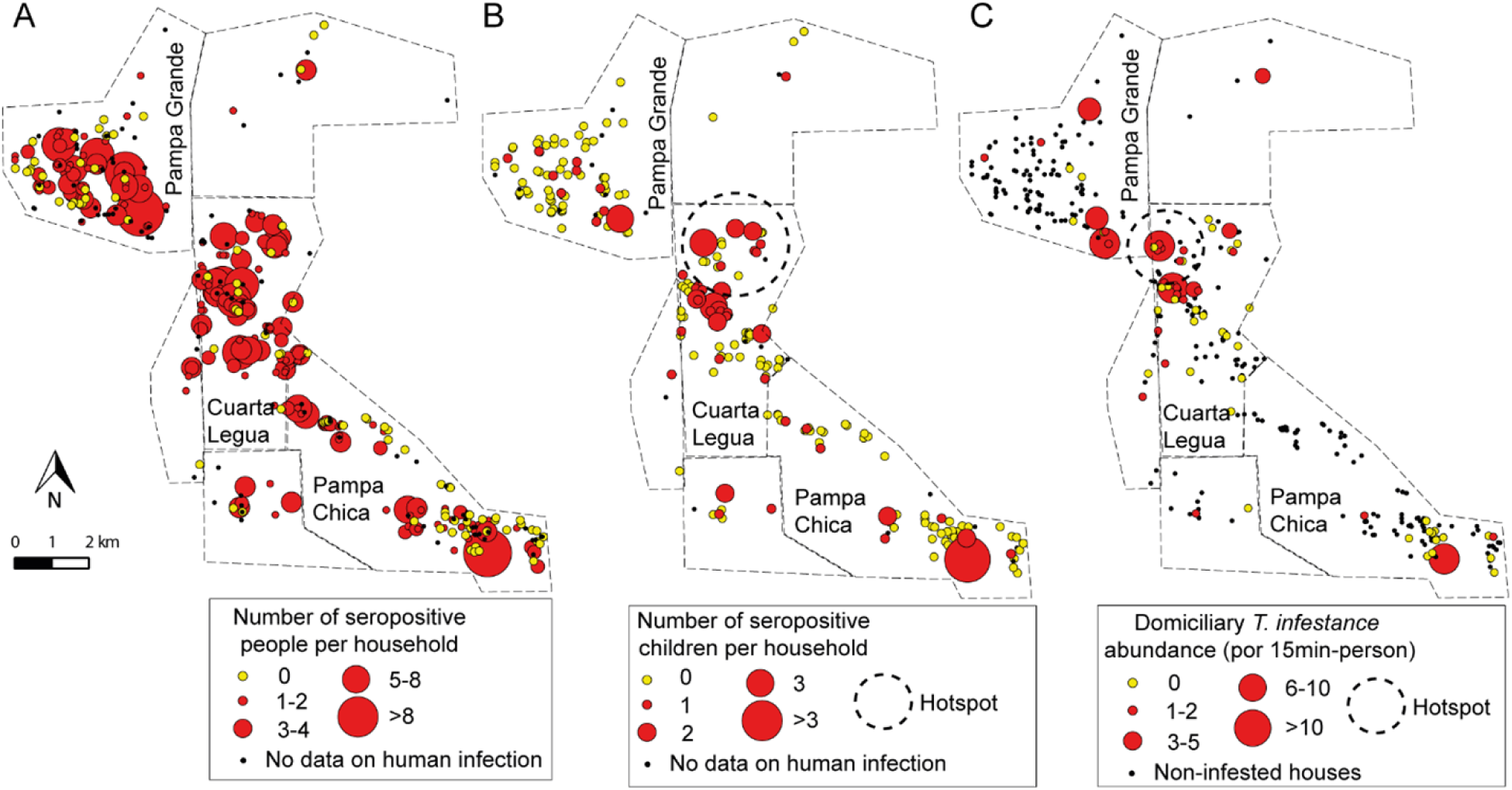
Distribution of the household number of *T. cruzi*-seropositive people **(A)** and children **(B)**, and the relative abundance of infected *T. infestans* per unit of effort **(C)** prior to community-wide insecticide spraying in 2008, Pampa del Indio, Argentina. The dashed circles represent the hotspot found by local spatial analysis. The maps were created in QGIS 2.18.11. based on the data collected within the scope of this study (S3 Table).

The proportion of houses with at least one seropositive child in 2008 also varied significantly between communities (χ^2^ test; df = 2; p < 0.01). The highest proportion was found in Cuarta Legua compared to the other communities (32.9% vs. 14.5-13.8%, respectively), but no spatial aggregation was found globally (S2C Fig). The number of infected children per household did not show a spatial correlation between neighboring households either (S2D Fig). However, local spatial analysis found a hotspot of child infection located in Cuarta Legua at a spatial scale between 1 and 2.6 km (Fig 5B). This hotspot was no longer detected in 2012-2015, nor did the spatial aggregation per community for the total population (S3 Fig).

The occurrence of at least one *T. cruzi*-infected *T. infestans* in the domicile was globally aggregated at all spatial scales when considering all houses (S2E Fig), but it did not differ from a random distribution when only infested houses were considered (S2F Fig). Nonetheless, a hotspot of domiciliary infected vectors was found in Cuarta Legua at a spatial scale between 0.2-1.8 km, which involved 6 houses with at least one infected vector (Fig 5C). This hotspot coincided partially with the child infection hotspot, but no significant correlation was found between human and vector infection patterns (S2G-H Fig).

The heatmaps show that the areas of high social vulnerability coincided with the qualitative and quantitative hotspots of human and vector infection (Fig 6). However, no significant spatial correlation was found between human or vector infection and socio-demographic variables (social vulnerability and host availability) (S4 Fig).

**Fig 6.**
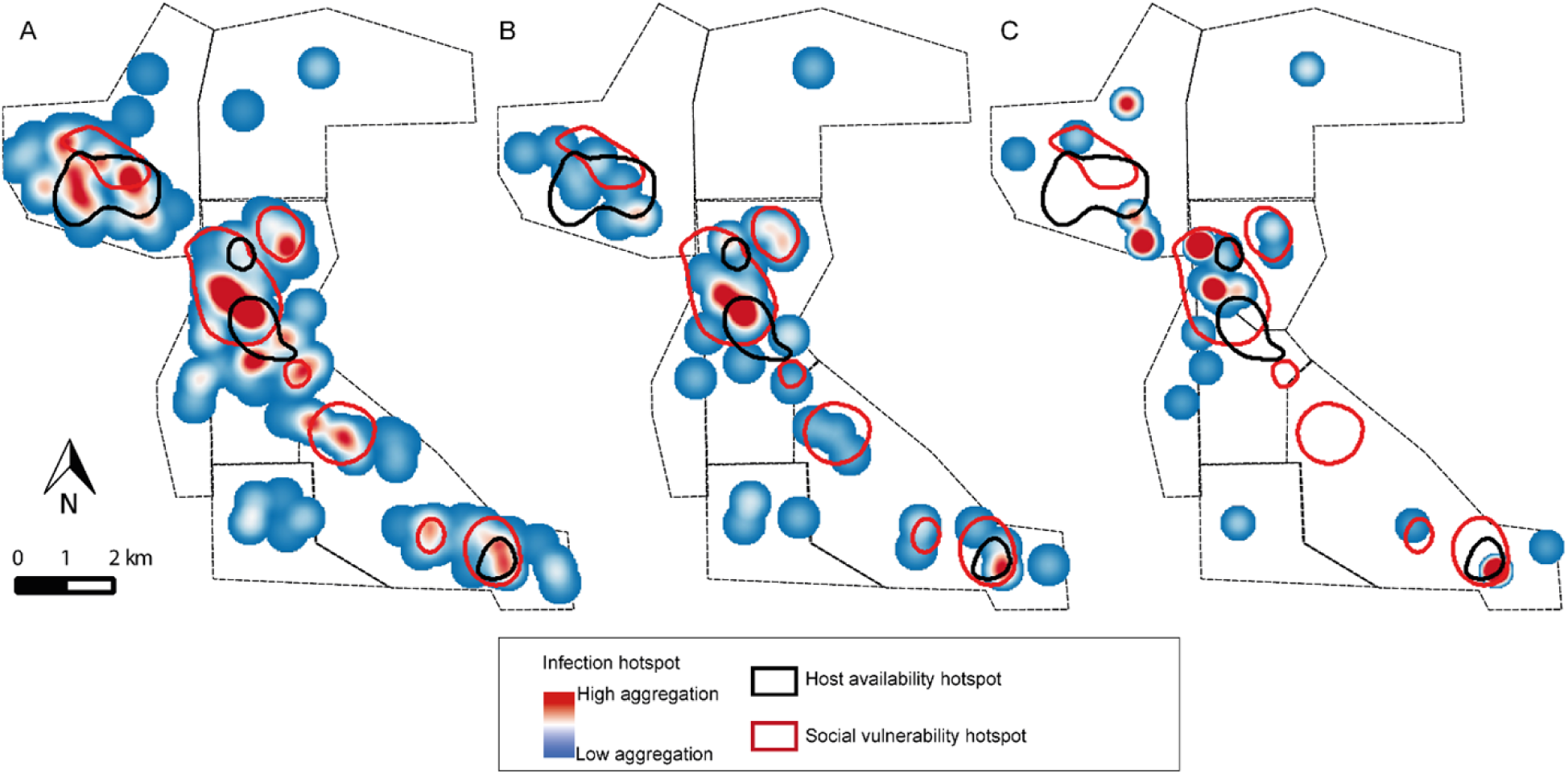
Heatmaps of the household number of *T. cruzi*-seropositive people **(A)** and children **(B)**, and the relative abundance of *T. cruzi*-infected *T. infestans* per unit of effort **(B)** in 2008, prior to a community-wide insecticide spraying campaign, Pampa del Indio, Argentina. The areas of qualitative aggregation (as determined by heatmaps) of social vulnerability and host availability are also shown. The maps were created in QGIS 2.18.11. based on the data collected within the scope of this study (S3 Table).

### Human infection risk model

Human infection was significantly clustered by household, after adjusting for the abundance of infected vectors, household social vulnerability and host availability (β_household_ = 1.3, CI_95_ = 1.02-1.7, log-likelihood ratio test, p < 0.001). When we included the number of seropositive co-inhabitants, the random effect of the household was nil and no significant differences were found between the GLMM and the respective GLM model (log-likelihood ratio test, p = 1), indicating that all variables included in the model accounted for the differences among households.

The risk of *T. cruzi* infection increased with age and if they were Qom, but did not vary by gender (Table 2). The ethnic group did not have a significant effect on child infection possibly because only one of the 70 seropositive children was creole. The risk of infection in the total population significantly increased by 60% for each additional infected *T. infestans* collected in the domicile and by 40% with each additional infected co-inhabitant; increased with increasing household social vulnerability, and decreased with increasing host availability index (Table 2).

**Table 2.** Generalized linear model (logit link function) of human seropositivity for *T. cruzi* infection for the total population (model 1) and children (model 2), Pampa del Indio, Argentina.

| *Model 1: population model* |  |  |  | *Model 2: children* |  |  |  |
| --- | --- | --- | --- | --- | --- | --- | --- |
| Variable | RI | OR (CI<sub>95</sub>) | P | Variable | RI | OR (CI<sub>95</sub>) | P |
| **Age (years)** | 1 | 1.07 (1.06-1.08) | <0.001** | **Age (years)** | 1 | 1.1 (1.04-1.3) | 0.008* |
| **Infected vector abundance (1)** | 1 | 1.6 (1.2-2.3) | 0.003* | **Infected vector abundance (1)** | 1 | 2.2 (1.001-4.8) | 0.04* |
| **Social vulnerability (2)** | 1 | 1.5 (1.2-1.8) | <0.001** | **Social vulnerability (2)** | 0.9 | 1.3 (0.8-2.1) | 0.2 |
| **Interaction (1) \* (2)** | 1 | 0.7 (0.6-0.9) | 0.01* | **Interaction (1) \* (2)** | 0.9 | 0.5 (0.2-0.98) | 0.04* |
| **Host availability** | 1 | 0.7 (0.6-0.9) | 0.003* | **Host availability** | 0.9 | 0.5 (0.2-0.9) | 0.02* |
| **Number of seropositive co-inhabitants** | 1 | 1.4 (1.3-1.6) | <0.001** | **Seropositive mother** | 0.9 |  |  |
| **Ethnic group** | 0.9 |  |  | No |  | 1 |  |
| Creole |  |  |  | Yes |  | 3.8 (1.4-11) | 0.009* |
| Qom |  | 2.3 (1.1-4.6) | 0.02* | **Gender** | 0.7 |  |  |
| **Gender** | 0.4 |  |  | Male |  | 1 |  |
| Male |  | 1 |  | Female |  | 2 (0.96-4.1) | 0.06~ |
| Female |  | 0.8 (0.6-1.1) | 0.2 | **Number of seropositive co-inhabitants** | 0.6 | 1.3 (0.96-1.9) | 0.08~ |
|  |  |  |  | **Recent insecticide spraying (2006-2008)** | 0.5 |  |  |
|  |  |  |  | No |  | 1 |  |
|  |  |  |  | Yes |  | 0.5 (0.2-1.3) | 0.2 |
|  |  |  |  | **Ethnic group** | 0.3 |  |  |
|  |  |  |  | Creole |  | 1 |  |
|  |  |  |  | Qom |  | 1.4 (0.2-12.5) | 0.7 |

\*\* $p < 0.001$ ; \* $0.001 \leq p \leq 0.05$ ; ~ $0.5 < p < 1$
OR: Odds ratio; RI: relative importance of the variable; CI<sub>95</sub>: 95% confidence interval

In the risk model for children, the positive effect of infected domestic vector abundance and the negative effect of host availability still held, but social vulnerability was no longer significant (Table 2). Including the mother’s seropositivity status (which was significantly and positively associated with child infection) led to a marginal effect of the number of seropositive co-inhabitants (Table 2). The recent partial insecticide spraying did not modify child infection risk (Table 2); it rather had a protective effect if the abundance of infected vectors was removed from the model (OR = 0.4, CI_95_ = 0.1-0.9, p = 0.03), indicating that recent insecticide treatment reduced the infection risk by decreasing vector abundance. In both models, we observed a significant negative interaction between social vulnerability and infected vector abundance; this indicates that the latter’s effect on infection risk was lower in households with higher social vulnerability (Table 2). Thus, people in households with high social vulnerability and low vector abundance have a similar infection risk as those in less vulnerable households but with higher vector abundance.

In both models, VIF<2 indicated no multicollinearity issues. The infection risk model for the total population had AUC = 0.83, with a sensitivity of 83% and a specificity of 72% (Fig 7). In a post-hoc classification, the model predicted 19.3% of false positives and 4.7% of false negatives. The model fit the data poorly according to the Hosmer-Lemeshow test (p < 0.001), perhaps due to the lack of saturation of the model, or non-linear relationship between variables, or limits to the goodness-of-fit test due to a large number of cases. By contrast, the risk model in children had a good fit to the data (Hosmer-Lemeshow test, p = 0.6); the AUC was 0.84 (Fig 7), and it had higher sensitivity (87%) and lower specificity (68%) than the previous model. It predicted 28.8% of false positive cases but only 1.1% of false negatives.

**Fig 7.**
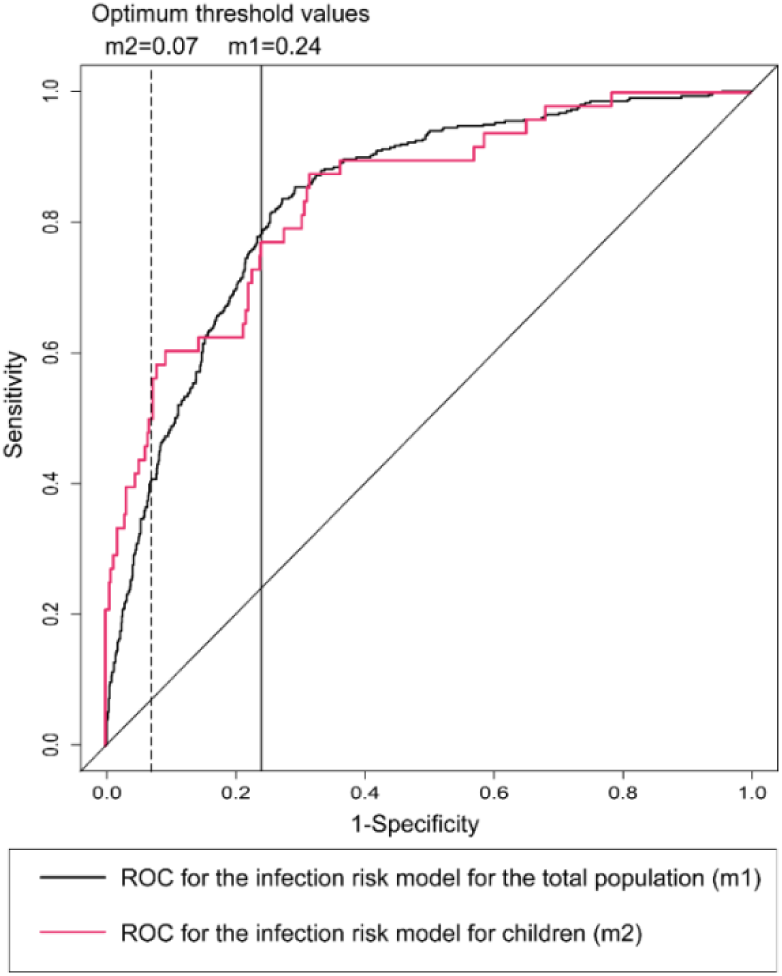
Receiver operating characteristic curves (ROC) of the *T. cruzi* infection risk models for the total population and children. The vertical lines show the optimum threshold value for classification (i.e. maximizes sensitivity and specificity).

The risk maps derived from the interpolation of the risk model documented the heterogeneous distribution of human infection and the occurrence of high-risk areas, which were more widespread when the total population was considered (Fig 8A) compared to the children risk map (Fig 8B). Although they failed to include a few households with seropositive children, most households with human cases were encompassed in these high-risk areas.

**Fig 8.**
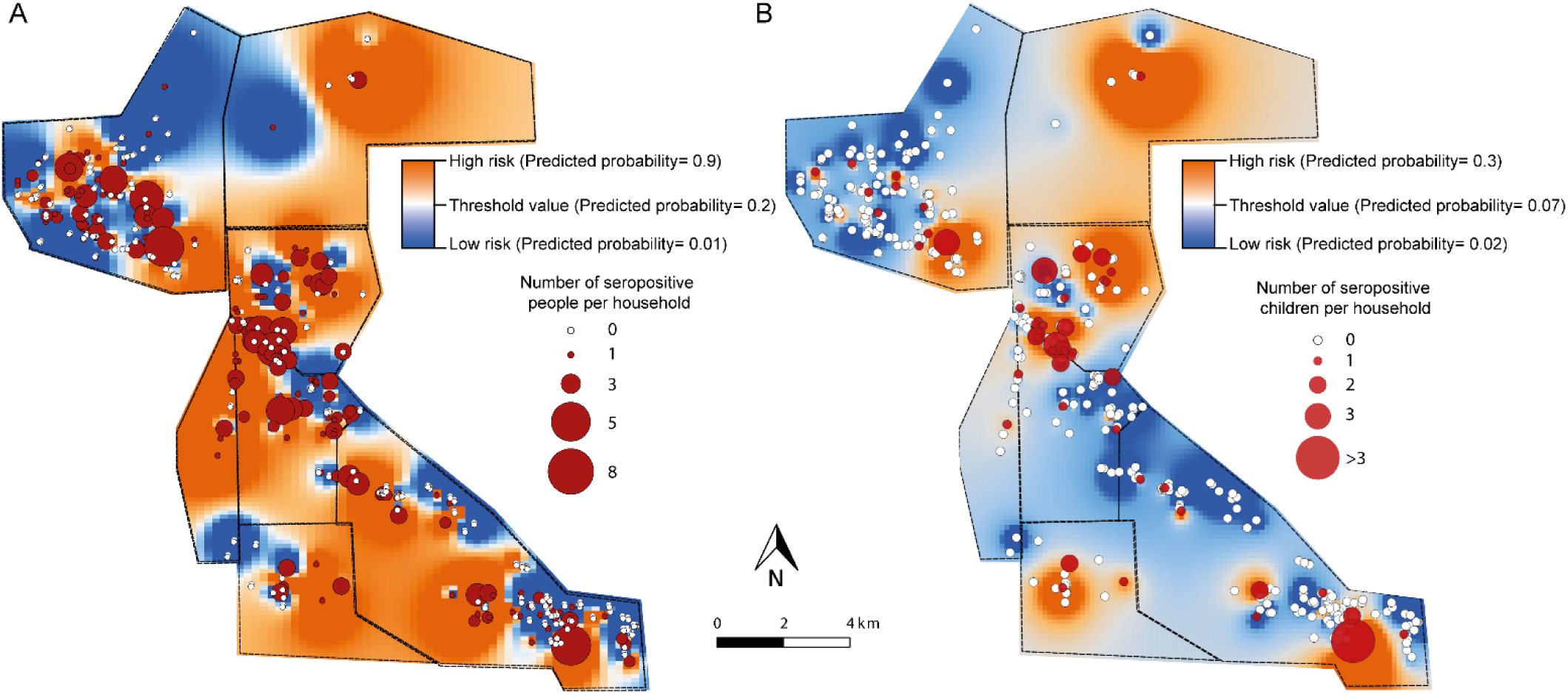
Risk maps of *T. cruzi* human infection for the total population **(A)** and children **(B)** in 2008, prior to community-wide insecticide spraying, Pampa del Indio, Argentina. The maps were created in QGIS 2.18.11. based on the data collected within the scope of this study (S3 Table).

## Discussion

This study shows the multivariate relationship among the ecological, demographic and socio-economic factors affecting human *T. cruzi* infection in endemic rural communities with active vector-borne transmission. To the best of our knowledge, this may be the first study to comprehensively assess their combined effects and interactions on human *T. cruzi* infection. In addition to corroborating the effects of demographic variables (age, ethnic group, among other) and the relative abundance of infected *T. infestans* on human infection risk [24], we found significant effects of socio-economic position after adjusting for ecological factors. Individual human infection risk significantly increased by 60% for each additional infected vector collected by unit of effort in the domicile and by 40% with each infected co-inhabitant; increased with increasing household social vulnerability, and decreased with increasing domestic host availability (numbers of indoor-resting bird or mammal hosts). A novel finding was the negative interaction between infected-vector abundance and social vulnerability, which indicates that members of more vulnerable households had an increased infection risk even at low infected-vector abundances. Household mobility also modulated infection risk: internal mobility (within the study area) reduced the effects of domiciliary vector abundance, possibly due to less consistent exposures among movers. Nonetheless, the seroprevalence rates of movers and non-movers were not significantly different. Although transmission was clustered by household, the spatial analysis showed that transmission risk was heterogeneous within the study area, with hotspots of human and vector infection matching areas of higher social vulnerability.

The Gran Chaco eco-region is a hotspot for Chagas disease and other NTDs that disproportionally affect vulnerable rural communities and certain demographic groups [11,65]. Like other NTDs, the highly focal distribution of Chagas disease cases is determined by particular combinations of various ecological and social determinants [65,66]. The baseline (2008) human seroprevalence here estimated (29%) was considerably lower than that registered in adjacent rural communities (40%) of Pampa del Indio [24]; in other indigenous (59-71%) and creole (40-62%) communities living in more disadvantaged, isolated areas of Chaco province known as “The Impenetrable” [17,18,21], and in the Bolivian Chaco (40-80%) [31,67]. Our seroprevalence estimate is similar to other estimates (28-31%) registered elsewhere in the Argentine Chaco [16,19]. This heterogenous distribution of human infection within Chaco province was evident in the 1980s when the intensity of vector-borne transmission was much higher [68], and remains evident almost three decades later.

The heterogeneous distribution of human infection was also captured within the studied communities through spatial analysis. Human cases, especially among children, were aggregated (Fig 5, S3 Fig) and matched the spatial aggregation of house infestation [41], vector infection and high social vulnerability (this paper). This pattern is consistent with vector-borne transmission mainly occurring at a household level [28,69], as human infection risk was mostly explained by household-level factors. Household clustering of cases was both revealed when we included the household as a random variable, and when the number of infected co-inhabitants was included in the human risk model. In the child infection model, household clustering was closely linked to the mother’s seropositivity status (allowing for putative congenital transmission), and to being under similar vector exposures [29,69].

The strong and positive association between human infection and the abundance of *T. cruzi*-infected vectors indicated active vector-borne transmission prior to the community-wide insecticide spraying we implemented. The occurrence of at least one seropositive child per household was significantly and positively associated with the abundance of infected *T. infestans* in the domicile, consistent with a more recent infection. Compared to the only previous study that quantified the effect of infected-vector domestic abundance on child infection risk, the magnitude we found was half of that estimated in rural communities of the dry Argentine Chaco two decades earlier (two-versus four-fold increase per each additional infected vector) [28]. However, the partial insecticide spraying undertaken within two years before the baseline vector survey, coupled with intense household mobility patterns [54] and slowly improving housing conditions [40], likely deflated the quantitative relationship between human and vector infection. Other studies reported a significant positive association between human infection and the presence of at least one infected domestic triatomine in rural communities [24,30,69], but not in a periurban community where child infection was instead associated with peridomestic infection in *T. infestans* [29]. In our study, because the housing units only had a few peridomestic sites [41], most of the infestations occurred in the domicile where human-vector contact with *T. infestans* is most likely to occur. Host availability is an ecological variable (i.e., number and density of host-feeding sources) closely related to household size. Here, we used a summary index to represent domestic host availability because usually there is a positive and significant correlation between household size and the number of domestic dogs or cats and chicken indoors [e.g., 54]. The occurrence of at least one seropositive child was associated positively with the domestic host availability index, as children were more likely to occur in larger households, and if other dimensions of risk are equal, there would be a greater chance of detecting at least one infected child in a larger household in which all members were screened. However, when the household number of infected (human) co-inhabitants was added to the individual risk model, the host availability index shifted from exerting positive effects to negative effects on human infection risk (weaker among children). Although there were no multicollinearity issues, understanding the effects of each type of host on the risk of human infection is hindered by the positive correlation between household size and the number of non-human domestic hosts, which pools key reservoir hosts (dogs and cats) with insusceptible hosts (chickens). Even though the host availability index exerted negative effects on the individual risk of infection, this result needs to be interpreted with caution since household size or domestic host abundance sometimes, but not always, were positively associated with larger domestic vector abundance [70,71]. The species identity of non-human hosts is indeed a key factor: one study found that child infection risk increased with the household number of dogs but decreased with the number of chicken indoors [28], whereas in another study human infection risk increased only with 2 or more infected dogs per household [24]. Unfortunately, our study lacks information on the distribution of *T. cruzi* infection in dogs and cats.

The household social vulnerability index used as a surrogate of socio-economic status represents a multidimensional measure of poverty, which we found associated with the domestic abundance of infected *T. infestans* [54]. Herein, we also found significant and positive effects of social vulnerability on human infection risk. To this aim we elaborated a comprehensive measure of socio-economic status; previous studies have focused on a few proxy variables: the goat-equivalent index [24], domicile’s characteristics (i.e., construction materials, size, cracks in walls, etc.) [28,31,32], and educational level [72]. The social vulnerability index summarized all of the above and also included household overcrowding, agricultural practices, at least one member with a stable source of income by employment, and the household number of welfare recipients [54]. The social vulnerability index also modified the effects of infected-vector abundance on human infection risk: the magnitude of this effect decreased as social vulnerability increased. This could be explained by several non-exclusive reasons: (i) the social vulnerability index in part may indicate a past, undetected vector infestation prior to the baseline survey in 2008 (i.e., past vector exposure); (ii) households with higher social vulnerability had greater local mobility [54], suggesting putative vector exposures elsewhere before the insecticide spraying campaign, and (iii) more frequent human-vector contacts given the smaller domicile and greater overcrowding in households with higher social vulnerability. Social vulnerability may also affect household resilience to face house infestation and the consequences of vector exposure. The access of vulnerable households to health services (for diagnosis and treatment, for example) was lower; they were less likely to use domestic insecticides [54] possibly due to economic constraints, and depended on government-sponsored vector control actions.

Household-level variables combined with ethnic group and age determined individual infection risk. In particular, ethnic group was a key demographic factor in stratifying infection risk; Qom people had a two-fold higher risk compared to creoles. Indigenous communities in the Argentine Chaco sometimes displayed higher infection risk than creole communities [17,21], but this pattern was not verified in rural communities in Pampa del Indio with a majority of creoles [24]. The effects of ethnic group on human infection, after adjusting for other social and ecological factors that co-vary with ethnicity, may be explained by differential mobility patterns between Qom and creole households [40,54]. The intense household mobility and the high rates of house turnover among the Qom most likely exerted negative impacts on *T. infestans* populations [41]. Movers were likely exposed to lower and more variable vector abundance over time, and also had higher social vulnerability and a higher risk of occupying an infested house at baseline (preintervention) [54,73]. Complex vector exposure patterns created by human mobility were reflected in the statistically significant interaction between domiciliary infestation and mobility in the child infection risk model. Although infection prevalence was not significantly different between movers residing in infested *versus* non-infested houses at baseline, domiciliary infestation had strong effects on non-movers’ infection risk. The predicted infection probability was highest for non-mover children in infested houses, indicating prolonged vector exposure. The impact of human mobility was also evident when the aggregated spatial distribution of human cases at baseline (2008) was compared with the random pattern recorded in 2012-2015. This particular trait of the Qom population needs to be considered when designing vector control actions and disease screening strategies, given that vector indices alone would only detect a fraction of the *T. cruzi*-seropositive children with internal mobility.

Levy et al. (2007) [29] proposed a two-step screening strategy for *T. cruzi*-infected children based on an individual infection risk model adapted to periurban areas of Arequipa (Peru) with a history of recent vector-borne transmission. Their best model included age, *T. infestans* abundance in the domicile, and presence of at least one infected vector. They proposed first to screen children by their age and domestic vector abundance, followed by screening children living in neighboring houses up to the distance of spatial aggregation [29]. The model proposed in this study had a higher diagnostic ability (AUC = 0.84 vs 0.72) and high sensitivity, which would allow to detect most of the infected children. Serological screening strategies that also include household socio-economic status and demographic characteristics, and if possible, the mother’s seropositivity status, are expected to perform better at the expense of added cost in obtaining such data. Such costs may be deflated by using the information for an integrated interventions across NTDs.

One limitation of this study was the temporal lag between the baseline vector survey (2008) and the human serosurveys (2012-2015), which required to retrospectively assign the examined individuals to their 2008 residential location. Thus, missing data for human infection increased as a result of considerable human mobility. Assignment errors were minimized by combining data from multiple sources and checking them with local primary healthcare agents, which led to a successful assignment of 80% of the population recorded in 2015. Human mobility patterns were only registered in 2012-2015; we assumed that this pattern was constant over time (i.e., movers in 2012-2015 also behaved as movers in 2008) based on the observation that movers did it more than once [54].

### Implications for disease control

The London Declaration Goals launched in 2012 proposed milestones to eliminate NTDs by 2020, including Chagas disease [74]. Access to diagnosis and treatment is one of the remaining challenges for sustainable control of Chagas disease in endemic areas [50,74]. Chagas disease national estimates, resonated by WHO [9], may fail to approximate the true magnitude of the problem [74]. Integrating the ecological and social determinants of human infection with the spatial component allowed us to identify individuals, households and geographic sectors at higher risk of infection. This information is key to design cost-effective serological screening strategies in resource-constrained areas for treatment of *T. cruzi*-infected people. Although our analysis took advantage of the detailed data collected during household surveys and census, the same approach could be done using national census data though probably at the coarser spatial resolution that the data would allow. Our infection risk model was also translated into an actionable tool (i.e., risk maps) that identified high-risk areas for local human infection and can point to higher-priority sections for targeted interventions oriented to suppress house (re)infestations, treat infected children, and thus reduce the burden of future disease. This approach may also be useful for integrated control interventions across NTDs.

## Acknowledgments

The authors thank the Chaco and National Chagas control programs for continuing field support and advice; M. Victoria Cardinal, Yael M. Provecho and Natalia P. Macchiaverna for contributions to fieldwork and questionnaire design; Fundación Mundo Sano for hospitality at the study site; local communities, healthcare workers and personnel from the local hospital “Dante Tardelli” for participating in the serosurveys and treatments, granting access to the laboratory, and especially to Drs. Nilda Blanco Acevedo, Omar Lana and Marcelo Wirth. The authors are also very grateful to the local communities for their warm welcome.

## Supporting information captions

**S1 Text.** Checklist of STROBE recommendations for observational studies.

**S1 Table.** Generalized linear mixed model of seropositivity for *T. cruzi* infection vs. demographic variables in 2008, clustered by household (logit link function).

**S2 Table.** Generalized linear mixed model of seropositivity for *T. cruzi* infection vs. demographic and vector indices in 2008, clustered by household (logit link function) for the total population and for children.

**S3 Table.** Individual-level and household-level database including data on human infection with *Trypanosoma cruzi*, domiciliary infestation with *Triatoma infestans*, and environmental and socio-demographic variables in Area III, Pampa del Indio.

**S1 Figure.** Number of people tested in each phase of the serosurvey in 2012-2015 in Area III, Pampa del Indio.

**S2 Figure.** Results from the global spatial point pattern analysis for the human and vector infection in 2008. (A) Univariate global analysis for the occurrence of at least one seropositive person in the household; (B) Variogram of the number of seropositive people per household; (C) Univariate global analysis for the occurrence of at least one seropositive children in the household; (D) Variogram of the number of seropositive children per household; (E) Univariate global analysis for the occurrence of at least one infected *T. infestans* considering all houses; (F) Univariate global analysis for the occurrence of at least one infected *T. infestans* considering only infested houses; (G) Bivariate global analysis for the correlation between children seropositivity and *T. infestans* infection; (H) Variogram for the spatial correlation between the abundance of infected *T. infestans* and the number of seropositive children per household. The lines with red dots indicate the observed values and the solid lines indicate the confidence ‘envelopes’.

**S3 Figure.** Distribution of the household number of *T. cruzi*-seropositive people **(A)** and children **(B)** in 2015. The maps were created in QGIS 2.18.11. based on the data collected within the scope of this study (S3 Table).

**S4 Figure.** Variograms from the global spatial point pattern analysis for socio-demographic variables and human or vector infection: the social vulnerability index vs. the number of seropositive people per household (A), vs. the number of seropositive children (B), and vs. the abundance of infected *T. infestans* (C); and the host availability index vs. the number of seropositive people per household (E), vs. the number of seropositive children (F), and vs. the abundance of infected *T. infestans* (G). The lines with red dots indicate the observed values and the solid lines indicate the confidence ‘envelopes’.

